# Accelerating Single-Cell Sequencing Data Analysis with SciDAP: A User-Friendly Approach

**DOI:** 10.1101/2024.02.28.582604

**Authors:** Michael Kotliar, Andrey Kartashov, Artem Barski

**Affiliations:** Division of Allergy and Immunology, Cincinnati Children’s Hospital Medical Center, Cincinnati, OH, 45229, USA; Division of Human Genetics, Cincinnati Children’s Hospital Medical Center, Cincinnati, OH, 45229, USA; University of Cincinnati College of Medicine, Cincinnati, OH, 45267, USA; Datirium, LLC, Cincinnati, OH, USA

**Keywords:** Single-cell, scRNA-Seq, scATAC-Seq, scMultiome, CWL, SciDAP

## Abstract

Single-cell (sc) RNA, ATAC and Multiome sequencing became powerful tools for uncovering biological and disease mechanisms. Unfortunately, manual analysis of sc data presents multiple challenges due to large data volumes and complexity of configuration parameters. This complexity, as well as not being able to reproduce a computational environment, affects the reproducibility of analysis results. The Scientific Data Analysis Platform (https://SciDAP.com) allows biologists without computational expertise to analyze sequencing-based data using portable and reproducible pipelines written in Common Workflow Language (CWL). Our suite of computational pipelines addresses the most common needs in scRNA-Seq, scATAC-Seq and scMultiome data analysis. When executed on SciDAP, it offers a user-friendly alternative to manual data processing, eliminating the need for coding expertise. In this protocol, we describe the use of SciDAP to analyze scMultiome data. Similar approaches can be used for analysis of scRNA-Seq, scATAC-Seq and scVDJ-Seq datasets.

## Introduction

Over the last decade, single-cell (sc) sequencing data analysis has become pivotal for uncovering novel biological and disease mechanisms. In one of the recent studies [1], it helped to reveal the signature of tumor-reactive T cells in human ovarian cancer. As the approaches used in this manuscript can be applicable to other studies, the authors took pains to carefully document all the steps of the analysis in a separate protocol paper [2]. However, even with the detailed methods section describing sample preparation and data analysis [2], one cannot guarantee reproducibility, especially when it comes to manual processing of the collected data. This lack of exact reproducibility can be due to several reasons, such as inconsistency of computational environment. Over time, the software used in the original analysis may become outdated or sometimes even deprecated.

Replacing it with the latest available version may lead to compatibility issues and discrepancies in the analysis results. Additionally, manual processing requires programming skills and is prone to human error. Use of a specialized analysis platform such as SciDAP (https://SciDAP.com) overcomes these problems [3]. The platform not only guarantees the reproducibility of the computational studies, but also provides an intuitive web-based interface (Fig. 1), eliminating the need for coding expertise.

**Fig. 1.**
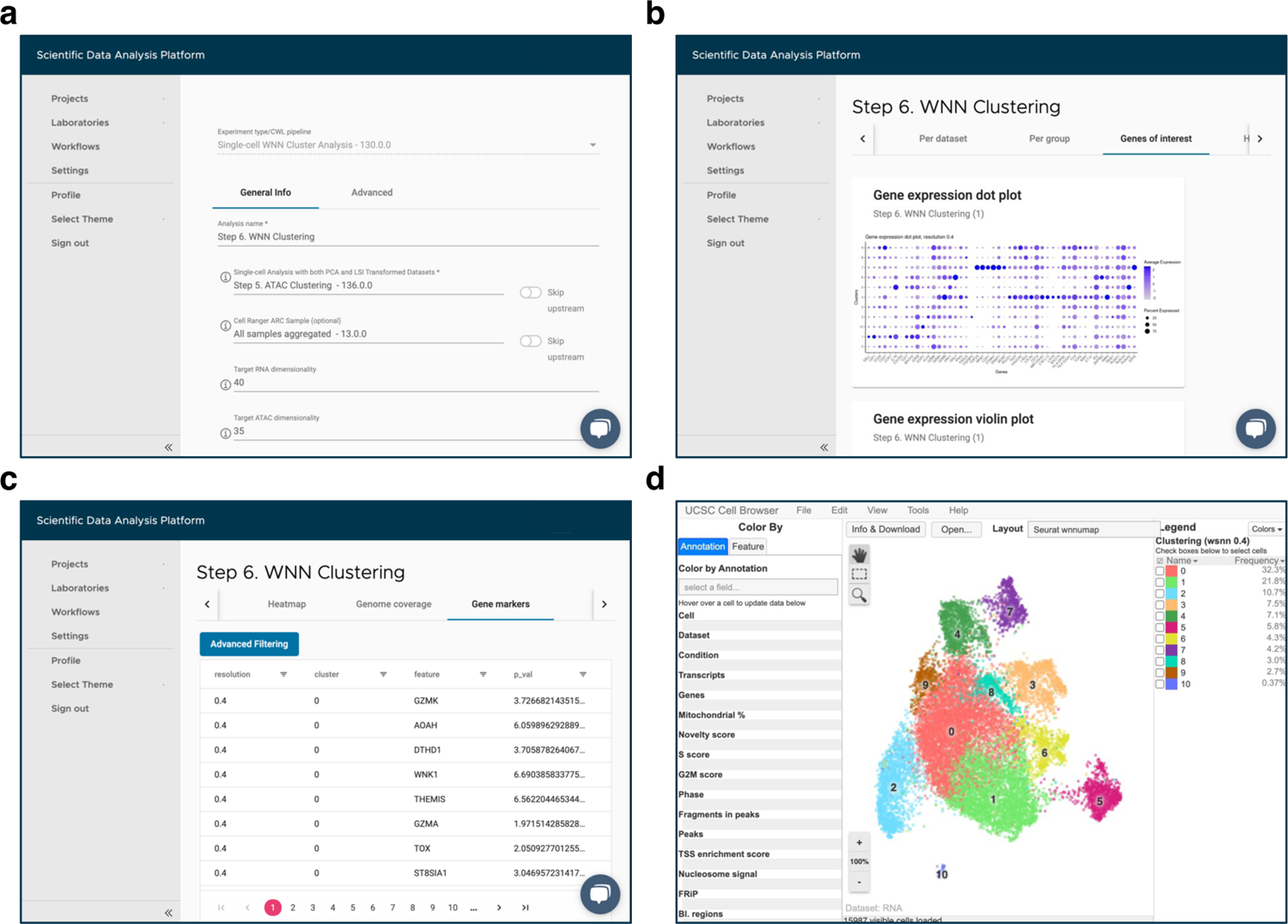
SciDAP web-based interface for adding a new analysis and viewing the experiment results. (a) Input parameters form for the “Single-Cell WNN Cluster Analysis” pipeline. (b) Expression of the user-provided genes on the “Genes of interest” tab. (c) Table with identified gene markers on the “Gene markers” tab. (d) Interactive visualization of the experiment results in built-in UCSC Cell Browser

All data analysis workflows are containerized and wrapped into CWL [4] (Common Workflow Language) format to guarantee independence from computational environment, reproducibility and portability. Therefore, exactly the same software version is run no matter how much time has passed since the original analysis had been developed. This is particularly important for ongoing, long-term studies, where the new datasets should be analyzed in exactly the same manner as the previous ones. Altogether, SciDAP helps biologist focus on the biological questions relevant to their study rather than spending time on software installation and configuration problems. When it comes to the analysis of sc sequencing data, with its complexity and abundance of configuration parameters, the advantages of using a platform with a convenient user interface and standardized workflows become more apparent. Therefore, we have developed multiple pipelines (Table 1) to address the most common needs in sc data analysis, all of which are now available in SciDAP. To illustrate the benefits of using these pipelines over manual processing, we will reanalyze scMultiome sequencing data from the abovementioned study [1].

**Table 1.** Workflows for single-cell sequencing data analysis available on SciDAP.

| # | Workflow | Description |
| --- | --- | --- |
| 1 | Cell Ranger Reference<br>(RNA, ATAC,<br>RNA+ATAC) | Builds a reference genome of a selected species for quantifying gene expression and chromatin accessibility. The results of this workflow are used in all “Cell Ranger Count” and “Cell Ranger Aggregate” pipelines. |
| 2 | Cell Ranger Reference<br>(VDJ) | Builds a reference genome of a selected species for V(D)J contig assembly and clonotype calling. The results of this workflow are used in the “Cell Ranger Count (RNA+VDJ)” pipeline. |
| 3 | Cell Ranger Count<br>(RNA) | Quantifies single-cell gene expression of the sequencing data from a single 10x Genomics library. The results of this workflow are used in either the “Single-Cell RNA-Seq Filtering Analysis” or “Cell Ranger Aggregate (RNA, RNA+VDJ)” pipeline. |
| 4 | Cell Ranger Count<br>(ATAC) | Quantifies single-cell chromatin accessibility of the sequencing data from a single 10x Genomics library. The results of this workflow are used in either the “Single-Cell ATAC-Seq Filtering Analysis” or “Cell Ranger Aggregate (ATAC)” pipeline. |
| 5 | Cell Ranger Count<br>(RNA+ATAC) | Quantifies single-cell gene expression and chromatin accessibility of the sequencing data from a single 10x Genomics library in a combined manner. The results of this workflow are used in either the “Single-Cell Multiome ATAC-Seq |
|  |  | and RNA-Seq Filtering Analysis” or “Cell Ranger Aggregate (RNA+ATAC)” pipeline. |
| 6 | Cell Ranger Count (RNA+VDJ) | Quantifies single-cell gene expression and performs V(D)J contig assembly and clonotype calling of the sequencing data from a single 10x Genomics library in a combined manner. The results of this workflow are used in the “Single-Cell RNA-Seq Filtering Analysis”, “Single-Cell Immune Profiling Analysis”, or “Cell Ranger Aggregate (RNA, RNA+VDJ)” pipeline. |
| 7 | Cell Ranger Aggregate (RNA, RNA+VDJ) | Combines outputs from multiple runs of either the “Cell Ranger Count (RNA)” or “Cell Ranger Count (RNA+VDJ)” pipeline. The results of this workflow are used in the “Single-Cell RNA-Seq Filtering Analysis” and “Single-Cell Immune Profiling Analysis” pipelines. |
| 8 | Cell Ranger Aggregate (ATAC) | Combines outputs from multiple runs of the “Cell Ranger Count (ATAC)” pipeline. The results of this workflow are used in the “Single-Cell ATAC-Seq Filtering Analysis” pipeline. |
| 9 | Cell Ranger Aggregate (RNA+ATAC) | Combines outputs from multiple runs of the “Cell Ranger Count (RNA+ATAC)” pipeline. The results of this workflow are used in the “Single-Cell Multiome ATAC-Seq and RNA-Seq Filtering Analysis” pipeline. |
| 10 | Single-Cell RNA-Seq Filtering Analysis | Removes low-quality cells from the outputs of the “Cell Ranger Count (RNA)”, “Cell Ranger Count (RNA+VDJ)”, and “Cell Ranger Aggregate (RNA, RNA+VDJ)” pipelines. The results of this workflow are used in the “Single-Cell RNA-Seq Dimensionality Reduction Analysis” pipeline. |
| 11 | Single-Cell ATAC-Seq Filtering Analysis | Removes low-quality cells from the outputs of either the “Cell Ranger Count (ATAC)” or “Cell Ranger Aggregate (ATAC)” pipeline. The results of this workflow are used in the “Single-Cell ATAC-Seq Dimensionality Reduction Analysis” pipeline. |
| 12 | Single-Cell Multiome ATAC-Seq and RNA-Seq Filtering Analysis | Removes low-quality cells from the outputs of the “Cell Ranger Count (RNA+ATAC)” and “Cell Ranger Aggregate (RNA+ATAC)” pipelines. The results of this workflow are used in the “Single-Cell RNA-Seq Dimensionality Reduction Analysis” and “Single-Cell ATAC-Seq Dimensionality Reduction Analysis” pipelines. |
| 13 | Single-Cell RNA-Seq Dimensionality Reduction Analysis | Removes noise and confounding sources of variation by reducing dimensionality of gene expression data from the outputs of the “Single-Cell RNA-Seq Filtering Analysis” or “Single-Cell Multiome ATAC-Seq and RNA-Seq Filtering Analysis” pipeline. The results of this workflow are used in the “Single-Cell RNA-Seq Cluster Analysis” pipeline. |
| 14 | Single-Cell ATAC-Seq Dimensionality Reduction Analysis | Removes noise and confounding sources of variation by reducing dimensionality of chromatin accessibility data from the outputs of either the “Single-Cell ATAC-Seq Filtering Analysis” or “Single-Cell Multiome ATAC-Seq and RNA-Seq Filtering Analysis” pipeline. The results of this workflow are used in the “Single-Cell ATAC-Seq Cluster Analysis” pipeline. |
| 15 | Single-Cell RNA-Seq Cluster Analysis | Clusters cells by similarity of gene expression data from the outputs of the “Single-Cell RNA-Seq Dimensionality Reduction Analysis” pipeline. The results of this workflow are used in the “Single-Cell Manual Cell Type Assignment”, “Single-Cell RNA-Seq Differential Expression Analysis”, “Single-Cell RNA-Seq Trajectory Analysis”, and “Single-Cell Differential Abundance Analysis” pipelines. |
| 16 | Single-Cell ATAC-Seq Cluster Analysis | Clusters cells by similarity of chromatin accessibility data from the outputs of the “Single-Cell ATAC-Seq Dimensionality Reduction Analysis” pipeline. The results of this workflow are used in the “Single-Cell Manual Cell Type Assignment”, “Single-Cell ATAC-Seq Differential Accessibility Analysis”, and “Single-Cell ATAC-Seq Genome Coverage” pipelines. |
| 17 | Single-Cell WNN Cluster Analysis | Clusters cells by similarity on the basis of both gene expression and chromatin accessibility data from the outputs of the “Single-Cell RNA-Seq Dimensionality Reduction Analysis” and “Single-Cell ATAC-Seq Dimensionality Reduction Analysis” pipelines run sequentially. The results of this workflow are used in the “Single-Cell Manual Cell Type Assignment”, “Single-Cell RNA-Seq Differential Expression Analysis”, “Single-Cell RNA-Seq Trajectory Analysis”, “Single-Cell Differential Abundance Analysis”, “Single-Cell ATAC-Seq Differential Accessibility Analysis”, and “Single-Cell ATAC-Seq Genome Coverage” pipelines. |
| 18 | Single-Cell Manual Cell Type Assignment | Assigns identities to cells clustered with any of the “Single-Cell Cluster Analysis” pipelines. For “Single-Cell RNA-Seq Cluster Analysis” the results of this workflow are used in the “Single-Cell RNA-Seq Differential Expression Analysis”, “Single-Cell RNA-Seq Trajectory Analysis”, and—when combined with outputs from the “Cell Ranger Count (RNA+VDJ)” or “Cell Ranger Aggregate (RNA, RNA+VDJ)” workflow—in the “Single-Cell Immune Profiling Analysis” pipeline. For “Single-Cell ATAC-Seq Cluster Analysis”, the results of this workflow are used in the “Single-Cell ATAC-Seq Differential Accessibility Analysis” and “Single-Cell ATAC-Seq Genome Coverage” pipelines. For “Single-Cell WNN Cluster Analysis”, the results of this workflow are used in all of the above, except the “Single-Cell Immune Profiling Analysis” pipeline. |
| 19 | Single-Cell RNA-Seq Differential Expression Analysis | Identifies differentially expressed genes between any two groups of cells, optionally aggregating gene expression data from single-cell to pseudobulk form. |
| 20 | Single-Cell ATAC-Seq Differential Accessibility Analysis | Identifies differentially accessible regions between any two groups of cells, optionally aggregating chromatin accessibility data from single-cell to pseudobulk form. |
| 21 | Single-Cell RNA-Seq<br>Trajectory Analysis | Infers developmental trajectories and pseudotime from cells clustered by similarity of gene expression data. |
| 22 | Single-Cell ATAC-Seq<br>Genome Coverage | Generates genome coverage tracks from chromatin accessibility data of selected cells. |
| 23 | Single-Cell Differential<br>Abundance Analysis | Compares the composition of cell types between two tested conditions. |
| 24 | Single-Cell Immune<br>Profiling Analysis | Estimates clonotype diversity and dynamics from V(D)J sequencing data assembled into contigs. |
| 25 | FASTQ Download | Assists in downloading problematic single-cell sequencing data from NCBI SRA (National Center for Biotechnology Information Sequence Read Archive) repository. |

## Materials

For running sc sequencing data analyses on SciDAP, an active subscription or trial are required. Alternatively, the same open-source CWL pipelines, available from our GitHub repository [5], can be manually executed as command line tools by more experienced users for testing and developing purposes. To manually run our CWL pipelines a user will need Docker [6] and CWL-Airflow [7] or other CWL workflow runner installed. Minimum hardware requirements to manually execute all described data analysis steps sequentially: 128GB of RAM, 8 CPU, 4TB of storage. The source code for all our sc data analysis pipelines is written in R [8], containerized and wrapped in CWL format. Pipelines utilize the following R packages: Seurat [9], Signac [10], Harmony [11], scRepertoire [12], DESeq2 [13], Limma [14], Hopach [15], DAseq [16], Slingshot [17], EnhancedVolcano [18], ComplexHeatmap [19], Glimma [20], Circlize [21], DittoSeq [22], Nebulosa [23], scDblFinder [24], and Dynverse [25], as well as some Python packages, such as UCSC Cell Browser [26], MAnorm2 [27], and MACS2 [28]. Raw read alignment and cell calling is performed with 10x Genomics Cell Ranger 7.0.0 [29], Cell Ranger ATAC 2.1.0 [30], and Cell Ranger ARC 2.0.2 [31].

Video guides for using SciDAP for analysis of various data types are available on SciDAP Youtube channel: https://www.youtube.com/@scidapscientificdataanalys8786

## Methods

### Overview

The analysis of sc sequencing data includes several stages and can be roughly divided into preprocessing that assigns reads to cells and genes or peaks, low-quality cell removal, dimensionality reduction, clustering, and cell annotation steps [32, 33]. Each of these can be accomplished by running one or several pipelines designed for a specific sequencing data type (scRNA-Seq, scATAC-Seq or scMultiome). Most of our workflows are intended to be run sequentially, reusing the outputs of the previous step (Fig. 2).

**Fig. 2.**
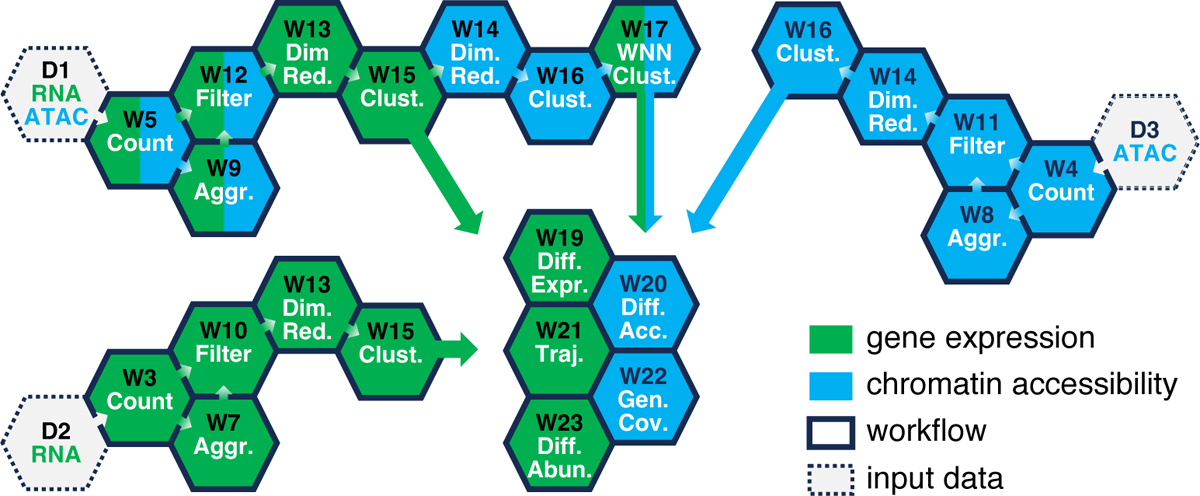
Workflow selection guide for analysis of scRNA-Seq, scATAC-Seq, or scMultiome data. Each hexagon labeled with “W” represents a workflow from Table 1. It can only be run when the upstream pipeline (the preceding hexagon linked by the arrow) has successfully completed. The color indicates the sequencing data type a pipeline can process. The W19-22 workflows can be run in parallel because they do not depend on each other. The workflow execution order begins with one of the hexagons labeled with “D” (representing input data) and progresses to adjacent hexagons following the arrows. A typical analysis of scMultiome data from multiple datasets begins with D1 and includes the W5→W9→W12→W13→W15→W14→W16→W17 and, optionally, W19-W22 steps.

Every pipeline produces interactive visualizations, tables, and publication-ready plots. Additionally, Seurat objects in the form R data files (.rds) are available for further analysis if needed. For scMultiome, the preprocessing step will include building the reference genome of a selected species, quantifying gene expression and chromatin accessibility (modalities) of the sequencing data from each of the datasets and merging the obtained results into a single feature-barcode matrix. The latter will include RNA reads per gene and ATAC fragments per peak counts for all cells found in the merged datasets. The next step is to identify and remove low-quality cells by jointly applying filtering thresholds to QC (quality control) metrics calculated for each modality. For scMultiome data, the main QC metrics include RNA reads, ATAC fragments, and genes per cell counts, percentage of RNA reads mapped to mitochondrial genes, TSS (Transcription Start Site) enrichment score, FRiP (Fraction of Reads in Peaks), nucleosome signal, and the fraction of ATAC fragments in genomic blacklist regions. Unlike low-quality cell removal, the dimensionality reduction of gene expression and chromatin accessibility data is performed independently by two different pipelines run sequentially. These steps normalize and integrate different experiments to ensure that similar cells from different samples cluster together. Each of these workflows also removes noise and possible confounding sources of variation by reducing the dimensionality in the selected modality. The next step is to cluster cells by similarity based on either gene expression or chromatin accessibility or both using WNN [34] (Weighted Nearest Neighbor) analysis. Resulting clusters usually recapitulate cell types or cell differentiation stages that can be annotated in a separate pipeline if needed. The detailed description of scMultiome data analysis in SciDAP is outlined below.

## 1 Creating a Project in SciDAP

In SciDAP, data analysis is organized into projects that contain both the data and the pipelines used to analyze them. After logging into SciDAP, click the “New project” button. On the left side of the window, select the “Overview” tab, and enter a title for the new project (e.g., “PRJNA793128”). You can optionally provide a subtitle and a description. On the right side of the window, go to the “Global” tab, and add the pipelines that we will use in this analysis by checking the boxes for the following workflows:

– FASTQ Download (optional, not needed if fastq files are already available)
– Cell Ranger Reference (RNA, ATAC, RNA+ATAC)
– Cell Ranger Count (RNA+ATAC)
– Cell Ranger Aggregate (RNA+ATAC)
– Single-Cell Multiome ATAC-Seq and RNA-Seq Filtering Analysis
– Single-Cell RNA-Seq Dimensionality Reduction Analysis
– Single-Cell ATAC-Seq Dimensionality Reduction Analysis
– Single-Cell RNA-Seq Cluster Analysis
– Single-Cell ATAC-Seq Cluster Analysis
– Single-Cell WNN Cluster Analysis
– Single-Cell Manual Cell Type Assignment

When done, click on “Save project”. If only scRNA-Seq or scATAC-Seq data are present, refer to Table 1 and Fig. 2 for selecting the proper workflows. For scRNA-Seq, begin with the D2 step and progress to the W3→W7→W10→W13→W15 steps, optionally including the W19, W21 and W23 pipelines. For scATAC-Seq, start from the D3 step and proceed to the W4→W8→W11→W14→W16 steps, optionally including the W20 and W22 pipelines.

## 2 Data Preprocessing

### 2.1 Downloading Single-Cell Sequencing Data

scMultiome analysis combines both gene expression and chromatin accessibility sequencing data; thus, it requires a minimum of five input FASTQ files per experiment. scRNA-Seq data are sequenced as paired-end, where R1 (at least 28 bases) contains a cell barcode and UMI (Unique Molecular Identifier) and R2 (at least 90 bases) is the cDNA sequence. The I1 and I2 (i7 and i5 indexes, respectively) files are used only for demultiplexing and are not needed for analysis. For scATAC-Seq, the i7 index file (I1) is not needed, but the i5 index file (I2, sometimes called R2, at least 24 bases) is required because it carries cell barcodes. The remaining two FASTQ files from scATAC-Seq (R1 and R2, sometimes called R1 and R3, at least 50 bases are recommended) contain DNA sequences of the Tn5 integration sites. For published studies, raw FASTQ files from scMultiome experiments are usually deposited to NCBI SRA (National Center for Biotechnology Information Sequence Read Archive) repository under two different SRR accession numbers for scRNA-Seq and scATAC-Seq data, respectively. Unfortunately, downloading raw FASTQ files from NCBI SRA is unnecessarily complicated because NCBI failed to establish and enforce a uniform way of storing the sc data, thus making programmatic access difficult [3]. Therefore, we recommend using the “FASTQ Download” pipeline described below. It can be omitted if FASTQ files are already available. The process of downloading sc FASTQ files for our example study is described below.

For each scMultiome experiment in Table 2, add two new samples (one with the accession number from the “RNA SRR” column, another from the “ATAC SRR” column) using the “FASTQ Download” pipeline.

**Table 2.** Sample metadata and SRR accession numbers.

| **Experiment name** | **Patient** | **Condition** | **RNA SRR** | **ATAC SRR** |
| --- | --- | --- | --- | --- |
| P_012320M_TRM | P_012320M | TRM | SRR17381636 | SRR17381635 |
| P_012320M_ReCir |  | ReCir | SRR17381634 | SRR17381633 |
| P_100809M_TRM | P_100809M | TRM | SRR17381632 | SRR17381631 |
| P_100809M_ReCir |  | ReCir | SRR17381630 | SRR17381629 |
| P_042718M_TRM | P_042718M | TRM | SRR17381628 | SRR17381627 |
| P_042718M_ReCir |  | ReCir | SRR17381626 | SRR17381625 |
| P_041218M_TRM | P_041218M | TRM | SRR17381624 | SRR17381623 |
| P_041218M_ReCir |  | ReCir | SRR17381622 | SRR17381621 |

To do this, in your project, click “Add sample”, and select “FASTQ Download” workflow from the dropdown menu at the top of the window. For “RNA SRR”, set the title for the new sample by copying it from the “Experiment name” column and adding the “Downloading RNA” prefix. For the “Comma or space separated list of SRR Identifiers” input, use the value from the “RNA SRR” column. Select “Split into all available files” from the “Split reads by” dropdown menu. Click on “Save sample”. Repeat a similar process for the accession number from the “ATAC SRR” column using the “Downloading ATAC” prefix in the sample’s title. Multiple samples can be added simultaneously. Downloaded FASTQ files are available in the “Files” tab of each sample that finished running successfully. An example of the outputs from the “Downloading ATAC P_041218M_ReCir” sample is shown in Fig. 3.

**Fig. 3.**
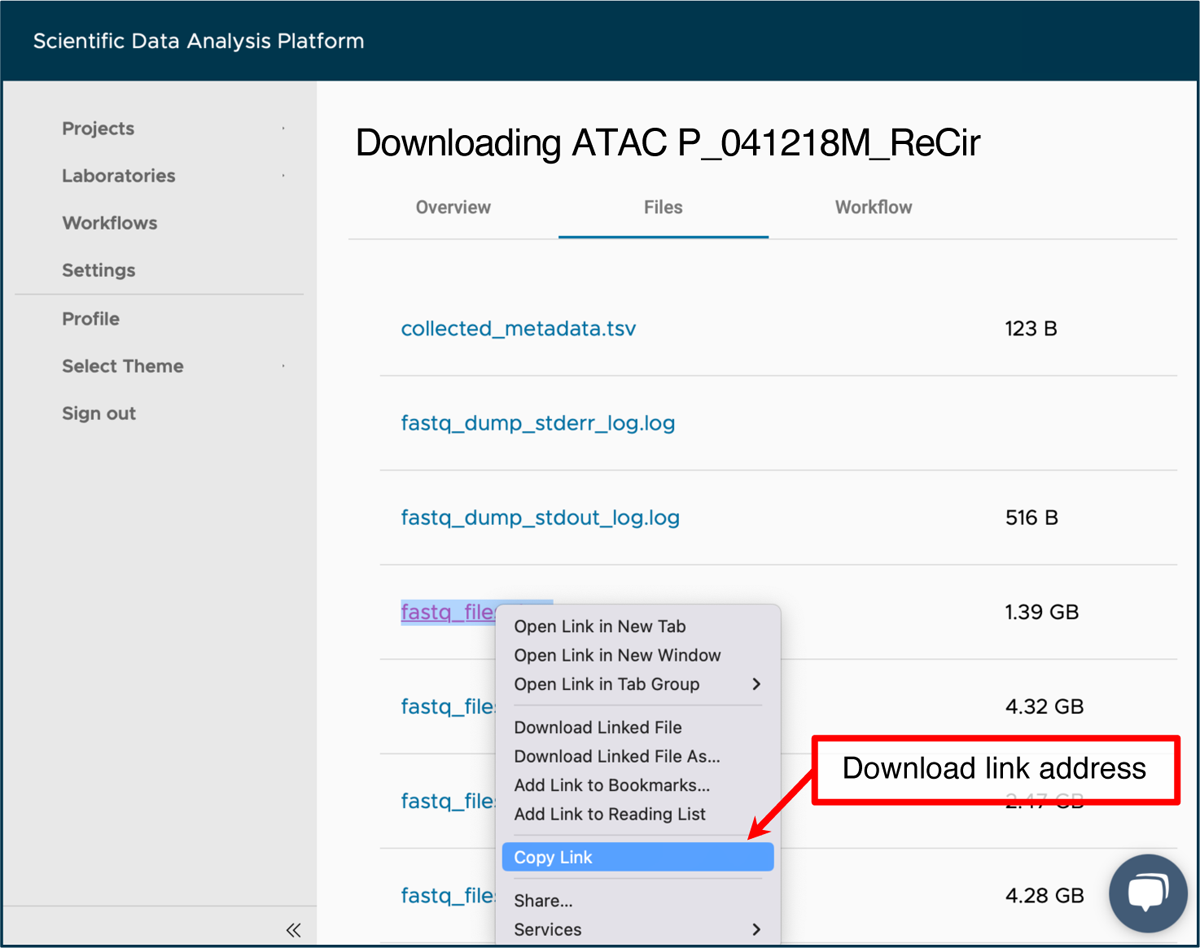
SciDAP web-based interface for accessing FASTQ files downloaded from the NCBI SRA repository by the “FASTQ Download” pipeline. When run with the “Split into all available files” parameter, the workflow will download and extract both technical and biological reads on the basis of the provided SRR accession number. The link address for files can be copied here for use in a later step.

### 2.2 Building a Reference Genome

Building genome indices is a required step before proceeding to gene expression and chromatin accessibility quantification, as the raw sequencing reads from the downloaded FASTQ files will be aligned to the reference genome using these indices. To do this, navigate to the “Sample” tab within the “PRJNA793128” project, click on “Add sample”, and choose the “Cell Ranger Reference (RNA, ATAC, RNA+ATAC)” workflow from the dropdown menu at the top of the window. In the form below, provide the title “Homo sapiens (hg38)” for a new experiment and set “Genome type” to “Homo sapiens (hg38)”. When done, click on “Save sample”. Wait for the experiment to finish running. If the same reference genome has been already built in another project, it can be shared with the current one. To share an already built reference genome a current project:

1. Navigate to the project with the existing reference genome sample.
2. Click on “Edit samples”.
3. Check the box near the sample with the reference genome.
4. From the dropdown menu below “Add sample” button, select the project with which to share the reference genome.
5. Click “Link samples”.
6. Click “Stop edit”.

### 2.3 Quantifying Single-Cell Gene Expression and Chromatin Accessibility

The next step is to quantify gene expression and chromatin accessibility for each cell. To do this, add a new sample using the “Cell Ranger Count (RNA+ATAC)” pipeline for each scMultiome experiment in Table 2 by following the instructions below.

1. Navigate to the “Sample” tab within the “PRJNA793128” project.
2. Open the “Downloading RNA [Experiment name]” sample for the current experiment.
3. In the “Files” tab, right click and copy the link addresses of the “fastq_files_2.gz” and “fastq_files_3.gz” outputs to be used for the “RNA FASTQ, Read 1” and “RNA FASTQ, Read 2” workflow inputs, respectively. The files can be identified by read length shown on the overview tab.
4. Navigate to the “Sample” tab within the “PRJNA793128” project.
5. Enter the “Downloading ATAC [Experiment name]” sample for the current experiment.
6. Copy the link addresses of the “fastq_files_2.gz”, “fastq_files_3.gz”, and “fastq_files_4.gz” outputs from the “Files” tab to be used for the “ATAC FASTQ, Read 1”, “ATAC FASTQ, Read 2”, and “ATAC FASTQ, Read 3” workflow inputs, respectively (Fig. 3).
7. Navigate to the “Sample” tab within the “PRJNA793128” project, and click on “Add sample”.
8. Select “Cell Ranger Count (RNA+ATAC)” workflow from the dropdown menu at the top of the window.
9. Provide a title for a new sample e.g. from the “Experiment name” column of Table 2.
10. Set the “Cell Ranger Reference Sample” to “Homo sapiens (hg38)”.
11. Attach all previously copied links to the appropriate workflow inputs by clicking on the “Use File Manager” button, selecting the “Attach From URL” tab, pasting a link into the designated field and clicking on the “Attach” button (Fig. 4b).

a. Alternatively, pre-downloaded FASTQ files can be uploaded to the “Current Sample” directory through the “File Manager” tab (Fig. 4a).
12. After all FASTQ files are attached to the respective inputs, click on “Save sample”. Multiple samples can be added simultaneously. Wait for all experiments to finish running.

**Fig. 4.**
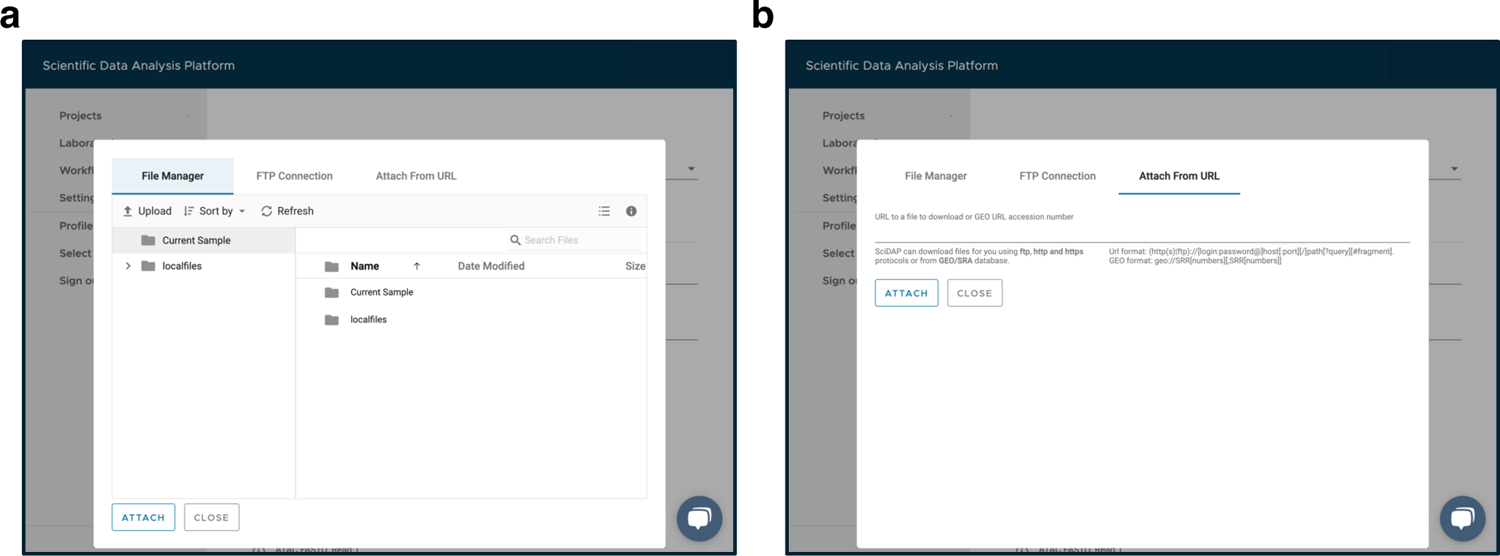
SciDAP web-based interface for file upload. (a) The “File Manager” tab for uploading local files to the platform. (b) The “Attach From URL” tab for adding a link address for a file to be uploaded to the platform.

### 2.4 Aggregating Outputs from Multiple Runs of the “Cell Ranger Count (RNA+ATAC)” Pipeline

To analyze cells from multiple datasets jointly, the results from all scMultiome experiments should be aggregated into a single feature-barcode matrix. To do this, navigate to the “Analysis” tab within the “PRJNA793128” project, click on “Add Analysis”, and choose the “Cell Ranger Aggregate (RNA+ATAC)” workflow from the dropdown menu at the top of the window. In the form below, provide a title “All samples aggregated” for the new analysis. Check the boxes next to all available samples in the “Cell Ranger RNA+ATAC Sample” dropdown menu. Before clicking on “Save sample”, switch to the “Advanced” tab and ensure that the “Library depth normalization” is set to “none” to prevent subsampling the reads to the smallest dataset. This subsampling may be useful for viewing the results with 10x Genomics Loupe Browser.

However, subsampling leads to data loss. Instead, normalization and scaling will be performed independently in scRNA-Seq and scATAC-Seq dimensionality reduction steps. Wait for the aggregation step to finish running.

## 3 QC Analysis and Low-Quality Cell Removal

### 3.1 Single-Cell Multiome ATAC-Seq and RNA-Seq Filtering Analysis

Removing low-quality cells is a crucial step in ensuring the accuracy and correct interpretation of scMultiome data analysis results. Filtering thresholds applied to multiple QC metrics help in selecting only high-quality cells. The thresholds proposed here are a good starting point and can be adjusted using the results of this analysis. When adjusting filtering thresholds, consider their joint effect, as some of the QC metrics correlate to each other. The easiest approach in this case is to examine distribution plots of the basic QC metrics and remove outlier peaks [35]. However, datasets containing heterogeneous cells may result in multiple peaks; therefore, select filtering thresholds to be as permissive as possible to avoid excluding viable cells. It is good practice to rerun the filtering step with updated parameters if clustering results show a strong batch effect related to any of the QC metrics. The filtering step can also be used for subsetting the data to selected cells for sub-clustering in the later steps by providing the list of cell barcodes. To remove low-quality cells in SciDAP, follow the instructions below.

1. Create a metadata table that will be used to group samples.

a. Navigate to the “Analysis” tab within the “PRJNA793128” project.
b. Enter the “All samples aggregated” experiment and download the “grouping_data.tsv” file from the “Files” tab. This file is a convenient template for the metadata table.
c. Open the downloaded file in a text editor, and edit the “condition” column to correspond to the values from the “Condition” column in Table 2. Ensure, that the order and number of rows in the edited file remain unchanged. These metadata (Table 3) will be used for highlighting cells on UMAP plots and differential analysis later. Additional metadata can be added at the later steps.
2. Navigate to the “Analysis” tab within the “PRJNA793128” project, click on “Add analysis”, and choose the “Single-Cell Multiome ATAC-Seq and RNA-Seq Filtering Analysis” workflow from the dropdown menu at the top of the window.
3. Provide the analysis name, e.g., “Step 1. Filtering”.
4. Select “All samples aggregated” from the dropdown menu below the “Cell Ranger RNA+ATAC Sample” label.
5. Click on “Use File Manager” under the “Datasets grouping (optional)” input, switch to the “File Manager” tab, enter the “Current Sample” directory, and upload previously prepared “grouping_data.tsv” file (Fig. 4a).
6. Select the uploaded file from the list, and click on the “Attach” button.
7. Switch to the “Advanced” tab and select “Based on either RNA or ATAC” in the “Doublets removal” dropdown menu. scDblFinder [24] will be used for doublet removal. This option is the strictest and will remove cells identified as doublets by either gene expression or chromatin accessibility data.
8. Update the “Minimum number of RNA reads per cell” and “Minimum number of genes per cell” inputs with 1000 and 500, respectively.
9. Set “Maximum mitochondrial percentage per cell” to 20%. This value is recommended for human cells [32]. Use 5% as a starting point for mouse cells.
10. Click on “Save sample”, and wait for the experiment to finish running.

**Table 3.**
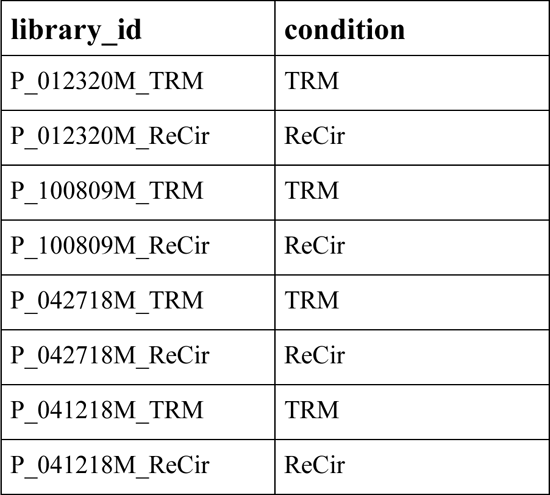
An example of the metadata file for splitting datasets into groups by the “condition” column.

Refer to Note 1 for a detailed description and default values of other configuration parameters. To access the analysis results enter the “Step 1. Filtering” experiment in the “Analysis” tab.

Choose the “Raw” or “Filtered” tab to view QC metrics plots before and after removing low-quality cells, respectively. For interactive exploration of the experiment results, select the “Overview” tab, and click on “UCSC Cell Browser”. All output files are also accessible for download on the “Files” tab. An example of the main QC metrics for a single dataset (P_100809M_ReCir) is shown in Fig. 5. After reviewing the results, the user may want to change some of the filtering parameters. To do so, the user can click on the “Edit” button within the sample, change the parameters and click on “Save sample” to re-run the filtering step.

**Fig. 5.**
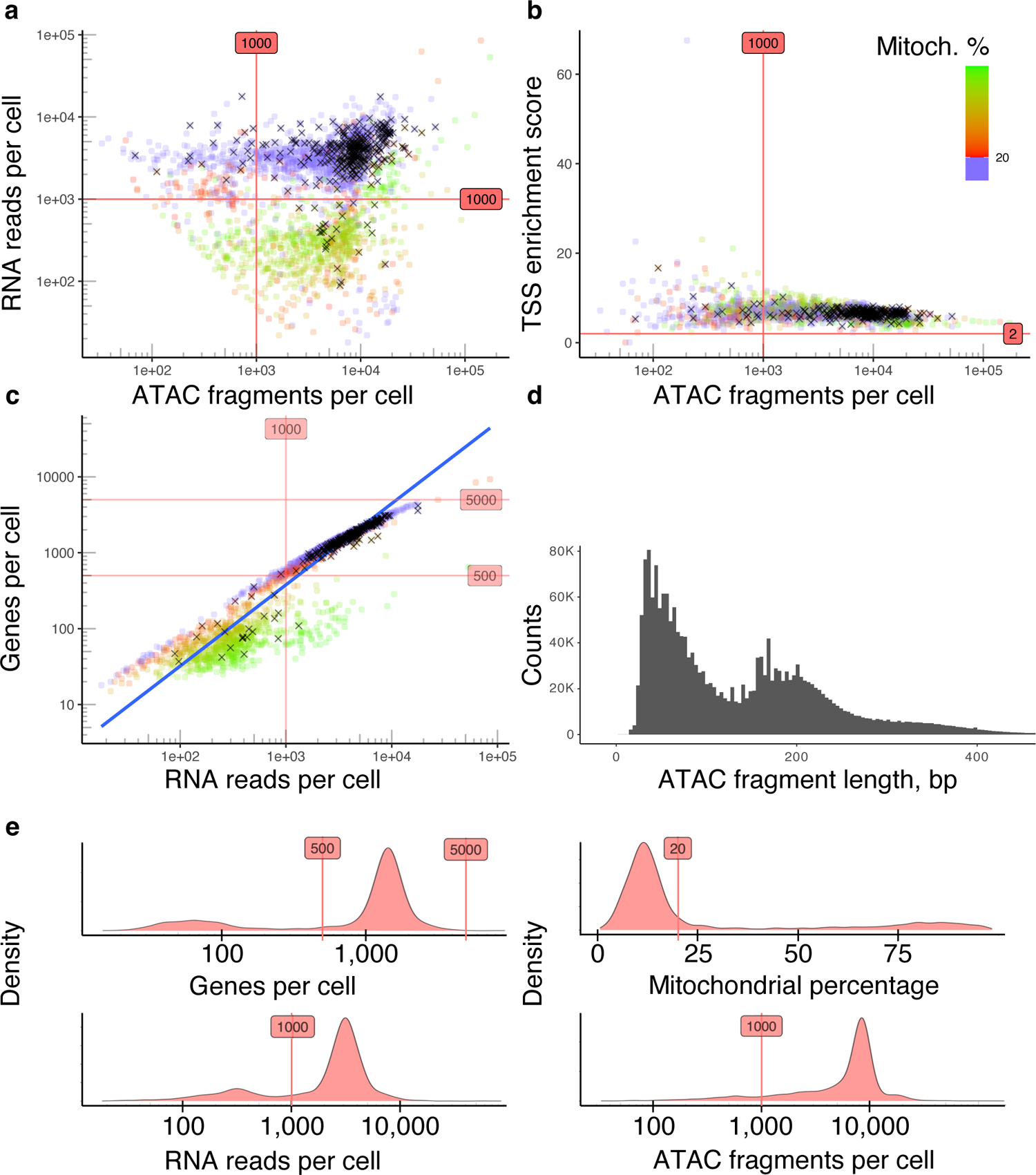
Assessing main QC metrics to remove low-quality cells from the P_100809M_ReCir dataset. (a) Include cells that have at least 1,000 ATAC fragments and 1,000 RNA reads per cell. Cells with a high fraction of mitochondrial reads (more than 20%) are discarded. (b) Include cells with a TSS enrichment score higher than 2. (c) Filter out cells with less than 500 or more than 5,000 genes, and exclude cells identified as doublets by either scRNA-Seq or scATAC-Seq data (marked with black crosses). (d) ATAC fragment length distribution for cells with a nucleosome signal below 4 (indicates high enrichment of fragments from nucleosome-free regions compared to the fragments with mono- and multi-nucleosomal lengths) (e) Density plots for evaluating main QC metrics and selected thresholds. The majority of viable cells (highest peaks) are not discarded.

## 4 Datasets Integration and Clustering

### 4.1 Single-Cell RNA-Seq Dimensionality Reduction Analysis

Not all of the genes are equally informative when it comes to grouping cells by their gene expression profiles. Moreover, high-dimensional sc data often contain noise and are prone to confounding sources of variation. Therefore, to capture the biologically meaningful signal, sc gene expression data usually undergo normalization, scaling, dimensionality reduction and, in case of multiple datasets merged together, integration procedures. In SciDAP, all these tasks can be performed by following the instructions below.

1. Create a metadata table that will be used to group samples.

a. Navigate to the “Analysis” tab within the “PRJNA793128” project.
b. Enter the “Step 1. Filtering” experiment and download the “datasets_metadata.tsv” file from the “Files” tab.
c. Open the downloaded file in a text editor and fill the “custom_patient” column with values from the “Patient” column in Table 2. Ensure, that the order and number of rows in the edited file remain unchanged. These metadata (Table 4) are used for assigning the datasets to groups for batch correction with Harmony.
2. Navigate to the “Analysis” tab within the “PRJNA793128” project, click on “Add analysis” and choose the “Single-Cell RNA-Seq Dimensionality Reduction Analysis” workflow from the dropdown menu at the top of the window.
3. Provide the analysis name “Step 2. RNA Dimensionality Reduction”.
4. Select “Step 1. Filtering” from the dropdown menu below the “Single-cell Analysis with Filtered RNA-Seq Datasets” label.
5. Select “harmony” in the “Integration method” dropdown menu and update the “Batch correction (harmony)” input with “custom_patient” value. This will remove patient-specific effects.
6. Set “Cell cycle gene set” to “human”.
7. Click on “Use File Manager” under the “Datasets metadata (optional)” input, switch to the “File Manager” tab, enter the “Current Sample” directory and upload the previously prepared “datasets_metadata.tsv” file (Fig. 4a).
8. Select the uploaded file from the list, and click on the “Attach” button.
9. Click on “Save sample”, and wait for the experiment to finish running.

**Table 4.** An example of the metadata file for dataset grouping.

| **library_id** | **custom_patient** |
| --- | --- |
| A P_012320M_TRM | P_012320M |
| B P_012320M_ReCir | P_012320M |
| C P_100809M_TRM | P_100809M |
| D P_100809M_ReCir | P_100809M |
| E P_042718M_TRM | P_042718M |
| F P_042718M_ReCir | P_042718M |
| G P_041218M_TRM | P_041218M |
| H P_041218M_ReCir | P_041218M |

Refer to Note 2 for a detailed description and default values of other configuration parameters. To access the analysis results enter the “Step 2. RNA Dimensionality Reduction” experiment in the “Analysis” tab. An elbow plot and other QC metrics, useful for quantifying datasets dimensionality and identifying potential confounding sources of variation, are shown on the “QC” tab. The quality of dataset integration can be evaluated through the plots available on the “Per dataset” and “Per group” tabs. For interactive exploration of the experiment results, select the “Overview” tab, and click on “UCSC Cell Browser”. All output files are also accessible for download on the “Files” tab. An example of 8 experiments integrated with Harmony is shown in Fig. 6.

**Fig. 6.**
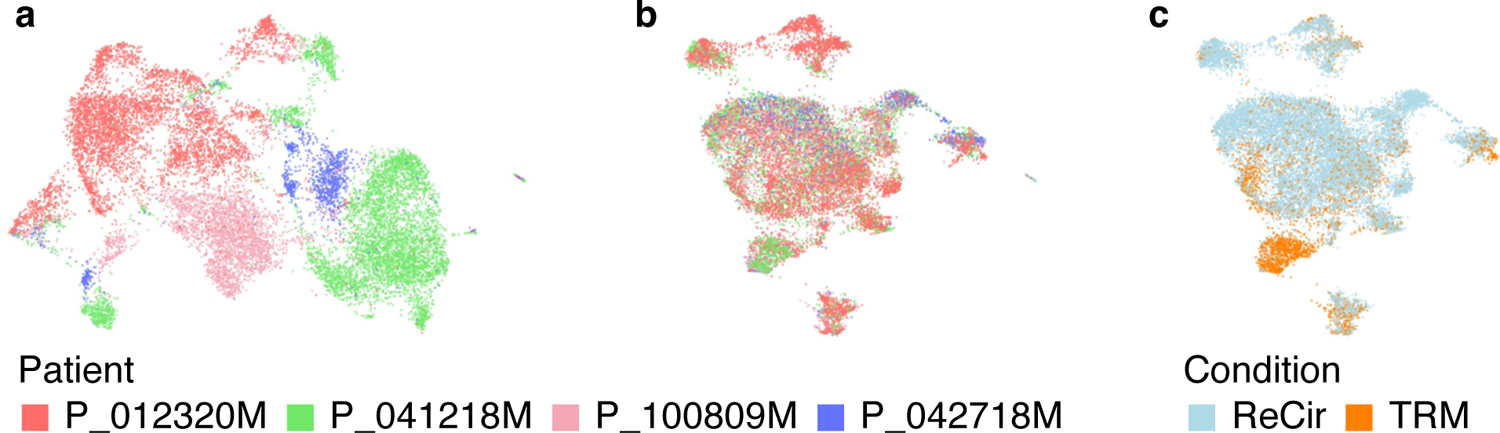
Evaluating the results of integrating scRNA-Seq data from 8 experiments. (a) The RNA UMAP plot from all datasets merged without integration. Cells from different patients form separate groups, which indicates the necessity of batch correction. (b, c) The RNA UMAP plots from the same datasets integrated with Harmony. The influence of the “custom_patient” covariate is removed, while preserving the expected distinction between TRM and recirculating CD8+ T cells.

### 4.2 Single-Cell RNA-Seq Cluster Analysis

The heterogeneity of sc gene expression data can be explored by clustering cells in the dimensionally reduced space. Resulting clusters usually resemble cell types or cell differentiation stages, each of which has a distinct set of gene markers (genes upregulated in the current cluster compared to all other cells). To run clustering analysis in SciDAP, follow the instructions below.

1. Navigate to the “Analysis” tab within the “PRJNA793128” project, click “Add analysis”, and select the “Single-Cell RNA-Seq Cluster Analysis” workflow.
2. Provide the analysis name “Step 3. RNA Clustering”.
3. Select “Step 2. RNA Dimensionality Reduction” from the dropdown menu below the “Single-cell Analysis with PCA Transformed RNA-Seq Datasets” label.
4. Update the “Clustering resolution” input with 0.5.
5. Copy the following genes into the “Genes of interest” input: “SELL, LEF1, CD28, CD27, CCR7, IL7R, CXCR5, TCF7, BACH2, JUNB, EGR1, KLF2, GZMK, GZMH, GZMB, PRF1, GNLY, IFNG, FASLG, FGFBP2, TOP2A, MKI67, CDK1, STMN1, DNMT1, MCM7, PDCD1, TIGIT, HAVCR2, LAG3, CTLA4, CD74, MIR155HG, CXCL13, LAYN, MYO7A, HLA-DRB1, HLA-DQA1, TOX, TOX2, BATF, ETV1, ID2, ZNF683, RBPJ, TBX21, RUNX1, RUNX3, RUNX2, EZH2”. This list does not affect the clustering but will be used to create several plots showing expression of these genes (e.g., Fig. 7b).
6. Click on “Save sample” button, and wait for the analysis to finish running.

**Fig. 7.**
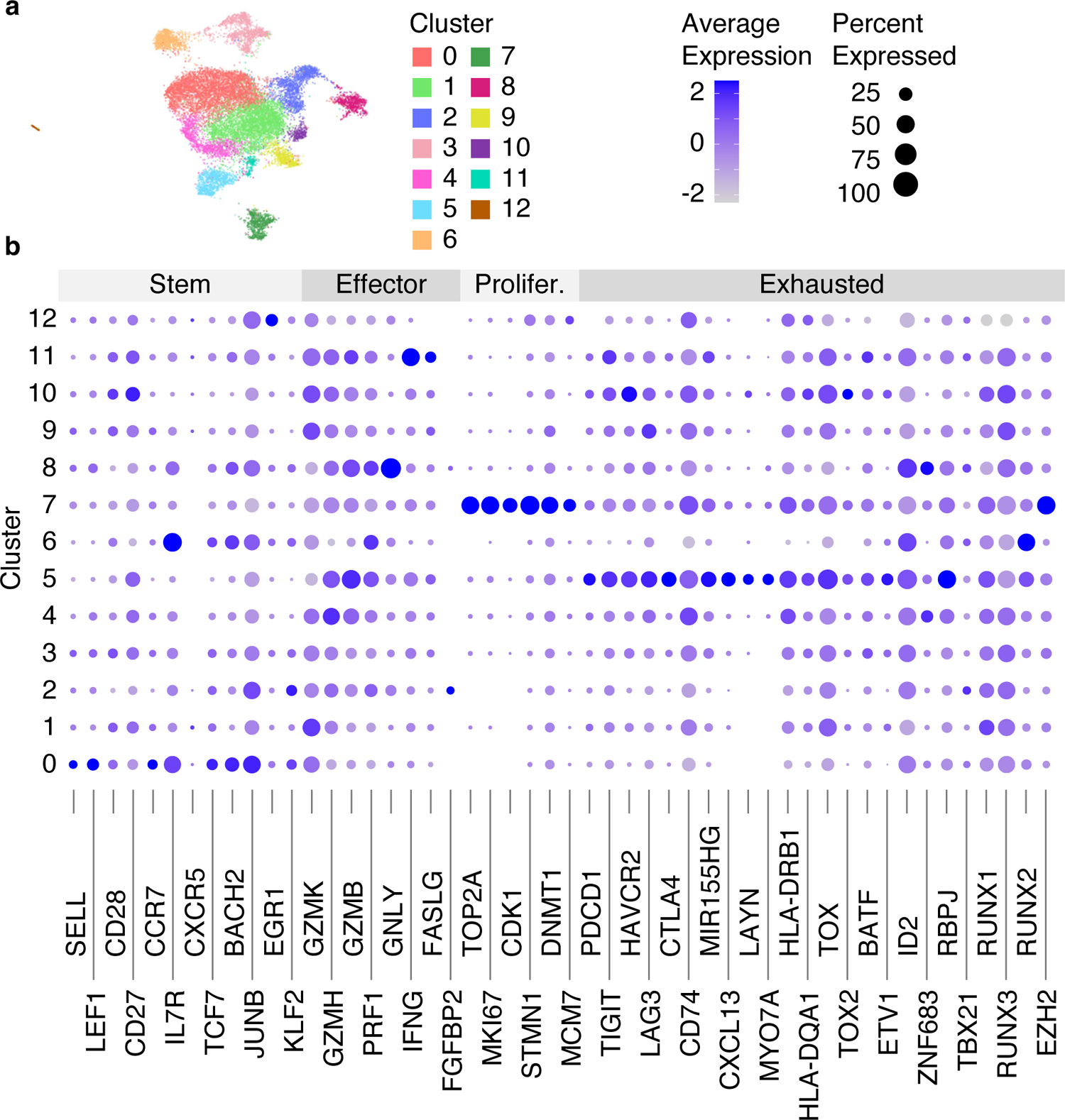
Clustering results of integrated with Harmony scRNA-Seq data from 8 datasets. (a) RNA UMAP plot with identified 13 clusters. (b) The dot plot showing the average gene expression and the percentage of cells per cluster for genes split into 4 categories (Stem, Effector, Proliferated [Prolifer.], Exhausted).

Refer to Note 3 for a detailed description and the default values of other configuration parameters. To access the analysis results enter the “Step 3. RNA Clustering” experiment in the “Analysis” tab. The RNA UMAP plot with identified clusters, as well as a silhouette score plot [36], used for evaluating the quality of clustering results, can be viewed on the “Per cluster” tab. The composition of each cluster categorized by dataset or grouping condition are shown on the “Per dataset” or “Per group” tabs, respectively. The expression of the user-provided genes can be explored through the plots available on the “Genes of interest” tab, while all gene markers are presented on the “Heatmap” and “Gene markers” tabs. For interactive exploration of the experiment results, select the “Overview” tab, and click on “UCSC Cell Browser”. All output files are also accessible for download on the “Files” tab. An example of clustering results is shown in Fig. 7.

### 4.3 Single-Cell ATAC-Seq Dimensionality Reduction Analysis

The feature-barcode matrix constructed from sc chromatin accessibility data shows number of reads per peak in each cell. Peaks can be called by Cell Ranger ARC 2.0.2 [31] based on pseudo-bulk of all cells or by MACS2 [28] separately for each cluster. To replace 10x peaks with the new ones, rerun “Single-Cell Multiome ATAC-Seq and RNA-Seq Filtering Analysis” step with the “Cell grouping for MACS2 peak calling” parameter set to the sc metadata column where the clusters are defined. Compared to scRNA-Seq, scATAC-Seq data are even more sparse because there are more open regions than genes. Moreover, each cell contains only 2 copies of DNA compared to multiple copies of mRNA. Representation of scATAC-Seq data in the dimensionally reduced space is often impacted by the sequencing depth; therefore, the first dimension (which is often highly correlated with the number of ATAC fragment counts in cell) is usually excluded from the analysis. Unlike scRNA-Seq data, integration of multiple scATAC-Seq datasets is typically performed through a shared low-dimensional space [9]. In SciDAP, all these tasks can be performed by following the instructions below.

1. Navigate to the “Analysis” tab within the “PRJNA793128” project, click on “Add analysis”, and choose the “Single-Cell ATAC-Seq Dimensionality Reduction Analysis” workflow from the dropdown menu at the top of the window.
2. Provide the analysis name “Step 4. ATAC Dimensionality Reduction”.
3. Select “Step 3. RNA Clustering” from the dropdown menu below the “Single-Cell Analysis with Filtered ATAC-Seq Datasets” label.
4. Set “Target dimensionality” to 35.
5. Click on “Save sample”, and wait for the experiment to finish running.

Refer to Note 4 for a detailed description and the default values of other configuration parameters. To access the analysis results enter the “Step 4. ATAC Dimensionality Reduction” experiment in the “Analysis” tab. Main QC metrics, useful for identifying potential confounding sources of variation, are shown on the “QC” tab. The quality of datasets integration can be evaluated through the plots available on the “Per dataset” and “Per group” tabs. For interactive exploration of the experiment results, select the “Overview” tab, and click on “UCSC Cell Browser”. All output files are also accessible for download on the “Files” tab. An example of 8 experiments integrated with Signac [10] is shown in Fig. 8.

**Fig. 8.**
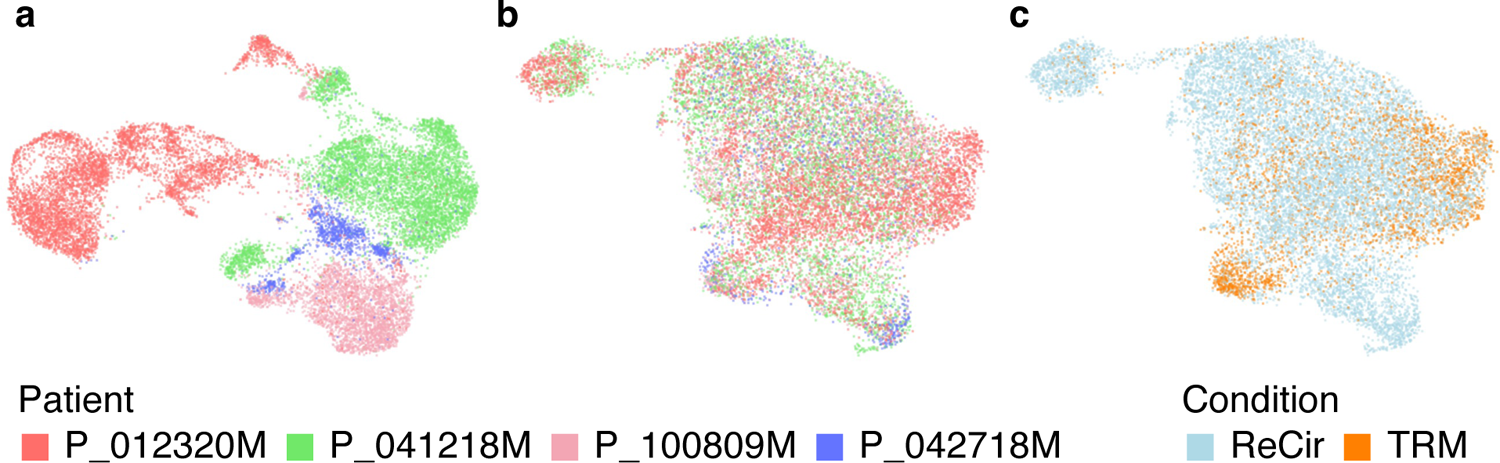
Evaluating the results of integrating scATAC-Seq data from 8 experiments. (a) The ATAC UMAP plot from all datasets merged without integration. Cells from different patients form separate groups, which indicates a noticeable batch effect. (b, c) The ATAC UMAP plots from the same datasets integrated with Signac. The technical differences between experiments are removed, while preserving the expected distinction between TRM and recirculating CD8+ T cells.

### 4.4 Single-Cell ATAC-Seq Cluster Analysis

Clustering of scATAC-Seq data is similar to scRNA-Seq; however, the obtained clusters reveal similarities in the cell regulatory states instead of common gene expression profiles. The first dimension is excluded from the analysis in the same manner as it was done in the “Single-Cell ATAC-Seq Dimensionality Reduction Analysis” step. To run clustering analysis in SciDAP, follow the instructions below.

1. Navigate to the “Analysis” tab within the “PRJNA793128” project, click on “Add analysis” and choose the “Single-Cell ATAC-Seq Cluster Analysis” workflow from the dropdown menu at the top of the window.
2. Provide the analysis name “Step 5. ATAC Clustering”.
3. Select “Step 4. ATAC Dimensionality Reduction” from the dropdown menu below the “Single-cell Analysis with LSI Transformed ATAC-Seq Datasets” label.
4. Select “All samples aggregated” from the dropdown menu below the “Cell Ranger ATAC or RNA+ATAC Sample (optional)” label.
5. Set “Target dimensionality” to 35.
6. Update the “Clustering resolution” input with 0.3.
7. Copy the following genes into the “Genes of interest” input: “TCF7, IL7R, PDGFB, GZMB, IFNG, TOX, ITGA2, ENTPD1, HAVCR2, EZH2”.
8. Click on “Save sample”, and wait for the experiment to finish running.

Refer to Note 5 for a detailed description and the default values of other configuration parameters. To access the analysis results enter the “Step 5. ATAC Clustering” experiment in the “Analysis” tab. The ATAC UMAP plot with identified clusters, as well as a silhouette score plot, used for evaluating the quality of clustering results, can be viewed on the “Per cluster” tab. The composition of each cluster categorized by dataset or grouping condition is shown on the “Per dataset” or “Per group” tabs, respectively. The ATAC fragment coverage around the user-provided genes can be explored through the plots available on the “Genome coverage” tab. For interactive exploration of the experiment results, select the “Overview” tab, and click on “UCSC Cell Browser”. All output files are also accessible for download on the “Files” tab. An example of clustering results is shown in Fig. 9.

**Fig. 9.**
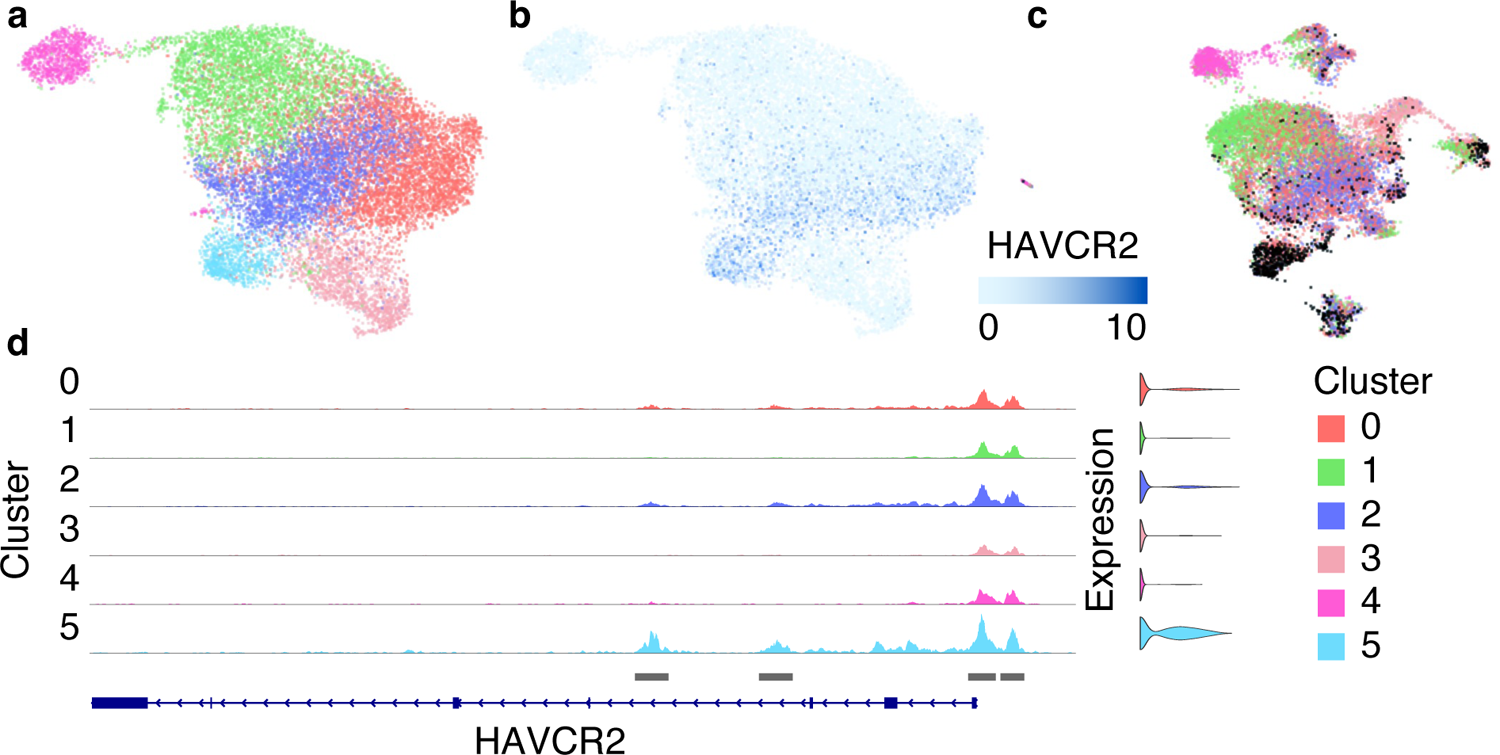
Clustering results of scATAC-Seq data from 8 datasets integrated with Signac. (a) The ATAC UMAP plot with identified 6 clusters. (b) The ATAC UMAP plot shows high expression of the exhaustion marker gene HAVCR2 in cluster 5. (c) The RNA UMAP plot colored by the clusters identified from scATAC-Seq data. Most cells from cluster 5 (marked with black) overlap with cluster 5 from the “Step 3. RNA Clustering” analysis. (d) ATAC fragment coverage plot around the HAVCR2 gene split by cluster.

### 4.5 Single-Cell WNN Cluster Analysis

The main advantage of scMultiome sequencing is capturing different types of information (modalities) from the same cells. Although gene expression and chromatin accessibility can be processed independently, as we demonstrated in the previous steps, an integrative analysis of both modalities can reveal novel information about the cell populations. WNN is one of the approaches that allows integration of multiple modalities and can handle the diverse information content within of each of them [34]. To perform WNN analysis in SciDAP, follow the instructions below.

1. Navigate to the “Analysis” tab within the “PRJNA793128” project, click on “Add analysis” and choose the “Single-Cell WNN Cluster Analysis” workflow from the dropdown menu at the top of the window.
2. Provide the analysis name “Step 6. WNN Clustering”.
3. Select “Step 5. ATAC Clustering” from the dropdown menu below the “Single-cell Analysis with both PCA and LSI Transformed Datasets” label.
4. Select “All samples aggregated” from the dropdown menu below the “Cell Ranger RNA+ATAC Sample (optional)” label.
5. Set “Target RNA dimensionality” to 40 and “Target ATAC dimensionality” to 35.
6. Update the “Clustering resolution” input with 0.4.
7. Enable the checkbox for the “Find gene markers” parameter.
8. Copy the following genes into the “Genes of interest” input: “SELL, LEF1, CD28, CD27, CCR7, IL7R, CXCR5, TCF7, BACH2, JUNB, EGR1, KLF2, GZMK, GZMH, GZMB, PRF1, GNLY, IFNG, FASLG, FGFBP2, TOP2A, MKI67, CDK1, STMN1, DNMT1, MCM7, PDCD1, TIGIT, HAVCR2, LAG3, CTLA4, CD74, MIR155HG, CXCL13, LAYN, MYO7A, HLA-DRB1, HLA-DQA1, TOX, TOX2, BATF, ETV1, ID2, ZNF683, RBPJ, TBX21, RUNX1, RUNX3, RUNX2, EZH2”.
9. Click on “Save sample”, and wait for the experiment to finish running.

Refer to Note 6 for a detailed description and default values of other configuration parameters. To access the analysis results enter “Step 6. WNN Clustering” experiment in the “Analysis” tab. WNN UMAP plot with identified clusters can be viewed on the “Per cluster” tab. The composition of each cluster categorized by dataset or grouping condition are shown on the “Per dataset” or “Per group” tabs, respectively. The expression of the user-provided genes can be explored through the plots available on the “Genes of interest” tab, while all gene markers are presented on the “Heatmap” and “Gene markers” tabs. The ATAC fragments coverage plots around the same genes of interest are available on the “Genome coverage” tab. For interactive exploration of the experiment results, select the “Overview” tab, and click on “UCSC Cell Browser”. All output files are also accessible for download on the “Files” tab. An example of integrated scRNA-Seq and scATAC-Seq data from 8 datasets using WNN analysis is shown in Fig. 10.

**Fig. 10.**
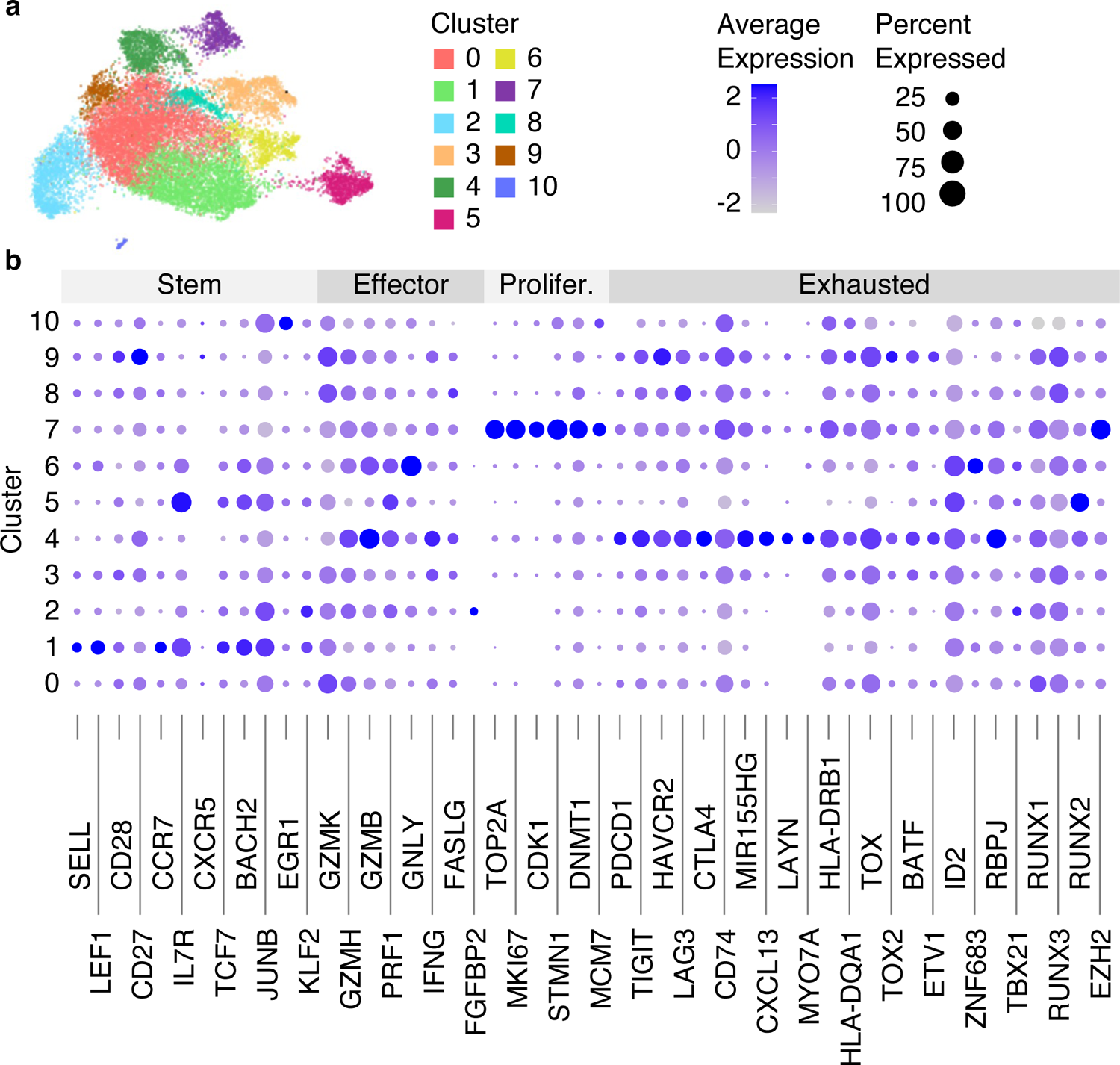
Clustering results of the integrated scRNA-Seq and scATAC-Seq data from 8 datasets using WNN analysis. (a) The WNN UMAP plot with identified 11 clusters. (b) The dot plot showing the average gene expression and the percentage of cells per cluster for genes split into 4 categories (Stem, Effector, Proliferated [Prolifer.], Exhausted).

## 5 Cell Annotation

### 5.1 Single-Cell Manual Cell Type Assignment

Cell annotation is the process of assigning identities to clusters on the basis of the expression of known marker genes. In scMultiome experiments, cell types are typically assigned on the basis of the clustering results of integrated modalities. It is also possible to use this pipeline to annotate clusters on the basis of scRNA-Seq or scATAC-Seq clustering instead. To manually annotate identified clusters from WNN analysis, follow the instructions below.

1. Using a text editor, create a comma-separated file “diff_stages.csv” with “cluster” and “celltype” columns.
2. Open the created file, and copy the cluster numbers from the results of the “Step 6. WNN Clustering” experiment to the “cluster” column.
3. Add values to the “celltype” column on the basis of the marker genes shown in Fig. 10b (e.g., “Stem” for clusters 1 and 5, “Effector” for cluster 6, “Proliferated” for cluster 7, “Exhausted” for cluster 4, and “Others” for all remaining clusters).
4. Navigate to the “Analysis” tab within the “PRJNA793128” project, click on “Add analysis” and choose the “Single-Cell Manual Cell Type Assignment” workflow from the dropdown menu at the top of the window.
5. Provide the analysis name “Step 7. Cell Differentiation Stages Assignment”.
6. Select “Step 6. WNN Clustering” from the dropdown menu below the “Single-cell Cluster Analysis” label.
7. Select “All samples aggregated” from the dropdown menu below the “Cell Ranger ATAC or RNA+ATAC Sample (optional)” label.
8. Select “WNN” in the “Dimensionality reduction” dropdown menu.
9. Update the “Clustering resolution” input with 0.4.
10. Copy the following genes into the “Genes of interest” input: “SELL, LEF1, CD28, CD27, CCR7, IL7R, CXCR5, TCF7, BACH2, JUNB, EGR1, KLF2, GZMK, GZMH, GZMB, PRF1, GNLY, IFNG, FASLG, FGFBP2, TOP2A, MKI67, CDK1, STMN1, DNMT1, MCM7, PDCD1, TIGIT, HAVCR2, LAG3, CTLA4, CD74, MIR155HG, CXCL13, LAYN, MYO7A, HLA-DRB1, HLA-DQA1, TOX, TOX2, BATF, ETV1, ID2, ZNF683, RBPJ, TBX21, RUNX1, RUNX3, RUNX2, EZH2”.
11. Click on “Use File Manager” under the “Cell types” input, switch to the “File Manager” tab, enter the “Current Sample” directory and upload the previously prepared “diff_stages.csv” file (Fig. 4a).
12. Select the uploaded file from the list, and click on the “Attach” button.
13. Click on “Save sample”, and wait for the experiment to finish running.

Refer to Note 7 for a detailed description and the default values of other configuration parameters. To access the analysis results enter “Step 7. Cell Differentiation Stages Assignment” experiment in the “Analysis” tab. The WNN UMAP plot with assigned cell differentiation stages can be viewed on the “Per cell type” tab. The composition of each cell differentiation stage categorized by dataset or grouping condition is shown on the “Per dataset” or “Per group” tabs, respectively. The expression of the user-provided genes can be explored through the plots available on the “Genes of interest” tab, while all gene markers are presented on the “Heatmap” and “Gene markers” tabs. The ATAC fragment coverage plots around the same genes of interest are available on the “Genome coverage” tab. If the “Criteria to split every cluster by (optional)” parameter is set to “condition” or “dataset”, each cell differentiation stage will be further split into the multiple groups showing each “condition” or “dataset” separately. For interactive exploration of the experiment results, select the “Overview” tab, and click on “UCSC Cell Browser”. All output files are also accessible for download on the “Files” tab. An example of the results from assigning different cell differentiation stages is shown in Fig. 11.

**Fig. 11.**
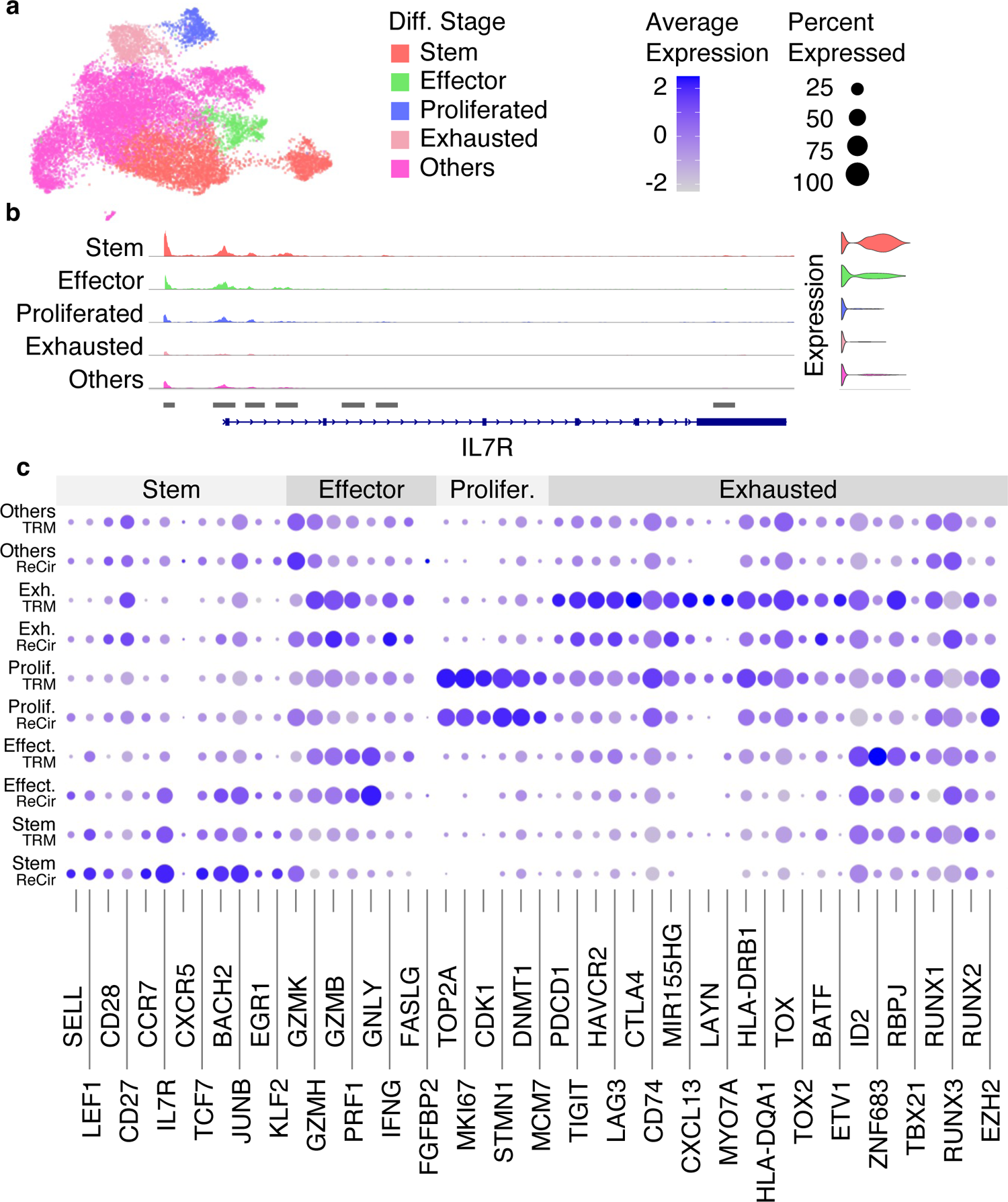
Results from assigning different cell differentiation stages. (a) The WNN UMAP plot with cells assigned to different differentiation stages (Diff. Stage). (b) The ATAC fragment coverage plot around the IL7R gene split by different cell differentiation stages. (c) The dot plot showing the average gene expression and the percentage of cells per cell type and per cell differentiation stage for genes split into 4 categories (Stem, Effector [Effect.], Proliferated [Prolifer.], Exhausted [Exh.]).

## 6 Additional Data Analysis Steps

The subsequent data analysis steps, not covered in this chapter, may be accomplished by running the “Single-Cell RNA-Seq Differential Expression Analysis”, “Single-Cell RNA-Seq Trajectory Analysis”, “Single-Cell Differential Abundance Analysis”, “Single-Cell ATAC-Seq Differential Accessibility Analysis”, and “Single-Cell ATAC-Seq Genome Coverage” pipelines (Table 1). Moreover, each of the W10-W17 and W21 workflows (Fig. 2) produce Seurat object with the analysis results saved in RDS (R Data Serialization) format.

## Notes

### 1 Single-Cell Multiome ATAC-Seq and RNA-Seq Filtering Analysis

If the experiment selected from the “Cell Ranger RNA+ATAC Sample” dropdown menu includes multiple aggregated datasets, each of them can be filtered independently by providing a comma- or space-separated list of filtering thresholds for the following parameters:

– Minimum number of RNA reads per cell
– Minimum number of genes per cell
– Maximum number of genes per cell
– Minimum novelty score per cell
– Minimum number of ATAC fragments in peaks per cell
– Minimum TSS enrichment score per cell
– Minimum FRiP per cell
– Maximum nucleosome signal per cell
– Maximum blacklist fraction per cell

The order and number of the specified values need to match the dataset order from the “aggregation_metadata.csv” output that is generated by selected “Cell Ranger RNA+ATAC Sample” and accessible on the “Files” tab.

#### Dataset grouping (optional)

If the selected “Cell Ranger RNA+ATAC Sample” includes multiple aggregated datasets, each dataset can be assigned to a separate group by providing a TSV/CSV file with “library_id” and “condition” columns (Table 3). Obtain a template of this file from the “grouping_data.tsv” output that is generated by “Cell Ranger RNA+ATAC Sample” and accessible on the “Files” tab.

#### Selected cell barcodes (optional)

Subsetting by cell barcodes can be used to identify sub-populations within the selected cluster(s) in the later steps. The user can obtain the list of barcodes by selecting cells in the UCSC Cell Browser. Cells of interest are identified by their barcodes and stored in a TSV/CSV file with at least one column named “barcode”. All other columns, except for “barcode”, will be added to the sc metadata loaded from “Cell Ranger RNA+ATAC Sample” and can be utilized in the current or future steps of analysis.

#### Cell grouping for MACS2 peak calling

The user can replace peaks called by Cell Ranger ARC 2.0 with the new ones. Peaks will be called by MACS2. There is a sc metadata column to group cells before running MACS2. To group cells by dataset, use “dataset”. Custom groups can be defined on the basis of any sc metadata column added through the “Selected cell barcodes (optional)” input. Default: use the original peaks generated by “Cell Ranger RNA+ATAC Sample”.

#### Minimum MACS2 FDR

There is a minimum FDR (q-value) cutoff for peaks identified by MACS2 if the “Cells grouping for MACS2 peak calling” input is provided. Default: 0.05

#### Doublet removal

There is a QC filtering parameter to remove cells identified as doublets by scDblFinder. Depending on the option selected, doublets can be detected and removed for gene expression or chromatin accessibility data singly or in combination (union or intersection). Default: “do not remove”.

#### Minimum number of RNA reads per cell

There is a QC filtering threshold to exclude from the analysis all cells with the number of RNA reads smaller than the provided value. Default: 500.

#### Minimum number of genes per cell

There is QC filtering threshold to exclude from the analysis all cells with the number of expressed genes smaller than the provided value. Default: 250.

#### Maximum number of genes per cell

There is a QC filtering threshold to exclude from the analysis all cells with the number of expressed genes larger than the provided value. Default: 5000.

#### Mitochondrial gene pattern

The Regex pattern can be used to identify mitochondrial genes on the basis of their names. Default: ^mt-|^MT-

#### Maximum mitochondrial percentage per cell

There is a QC filtering threshold to exclude from the analysis all cells with the percentage of RNA reads mapped to mitochondrial genes exceeding the provided value. Default: 5.

#### Minimum novelty score per cell

There is a QC filtering threshold to exclude from the analysis all cells with the novelty scores smaller than the provided value. This QC metric indicates the overall transcriptomic dissimilarity of the cells and is calculated as the ratio of log10(genes per cell) to log10(RNA reads per cell). Default: 0.8.

#### Minimum number of ATAC fragments in peaks per cell

There is a QC filtering threshold to exclude from the analysis all cells with the number of ATAC fragments in peaks smaller than the provided value. Default: 1000.

#### Minimum TSS enrichment score per cell

There is a QC filtering threshold to exclude from the analysis all cells with the TSS (Transcription Start Site) enrichment score smaller than the provided value. This QC metric is calculated on the basis of the ratio of ATAC fragments centered at the genes’ TSS to ATAC fragments in the TSS-flanking regions. Default: 2.

#### Minimum FRiP per cell

There is a QC filtering threshold to exclude from the analysis all cells with the FRiP (Fraction of Reads in Peaks) smaller than the provided value. Default: 0.15.

#### Maximum nucleosome signal per cell

There is a QC filtering threshold to exclude from the analysis all cells with the nucleosome signal higher than the provided value. The nucleosome signal is a measurement of nucleosome occupancy. It quantifies the approximate ratio of mononucleosomal to nucleosome-free ATAC fragments. Default: 4.

#### Maximum blacklist fraction per cell

There is a QC filtering threshold to exclude from the analysis all cells with the fraction of ATAC fragments in genomic blacklist regions larger than the provided value. Default: 0.05.

### 2 Single-Cell RNA-Seq Dimensionality Reduction Analysis

#### Normalization method

The normalization and scaling methods serve to remove technical variability between the cells: “sct” – uses SCTransform function for variance-stabilizing transformation [37]; “sctglm” – uses SCTransform function with glmGamPoi [38, 39] support for variance-stabilizing transformation; or “log” – uses NormalizeData and ScaleData functions for log-normalization and subsequent gene-level scaling [9]. All of these methods are preferred to the normalization by sub-sampling that is available in the “Cell Ranger Aggregate (RNA+ATAC)” and “Cell Ranger Aggregate (RNA, RNA+VDJ)” pipelines. When “log” is selected a single scaling factor will be applied across all genes in a cell. This leads to inefficient normalization of highly expressed genes [37]. Both “sct” and “sctglm” overcome this problem by using Pearson residuals from the regularized negative binomial regression model as the variance-stabilized gene expression levels. The latter are completely independent of the sequencing depth while preserving the biological variation. “sctglm” provides the support of the improved parameters estimation using glmGamPoi [39] R package that substantially increases the speed of calculation. Default: “sctglm”

#### Integration method

The integration methods serve to match shared cell types and states across experimental batches, donors, conditions or datasets: “seurat” – uses cross-dataset pairs of cells that are in a matched biological state (“anchors”) to correct for technical differences; “harmony” – uses Harmony algorithm to iteratively correct PCA (Principal Component Analysis) embeddings; or “none” – does not run integration, merges datasets instead. Select “harmony” when the batch effect is known and can be defined by the “Batch correction (harmony)” parameter. Compared to “seurat”, “harmony” requires less computational resources but might be biased toward major cell types [40]. Default: “seurat”

#### Batch correction (harmony)

When “harmony” is selected as “Integration method”, the batch effects are corrected on the basis of the provided factors. Specifically, “dataset” is used to integrate out the influence of the cells’ dataset of origin, while the factor “condition” is used to eliminate the influence of dataset grouping. Default: “dataset”

#### Target dimensionality

The target dimensionality is the number of principal components to be used in the PCA and UMAP projection, with accepted values ranging from 1 to 50. Use an elbow plot as a guide to identify the number of principal components where the curve starts to plateau. Default: 40.

#### Cell cycle gene set

Assign the cell cycle score and phase on the basis of the gene set for the selected organism. The available options are “human”, “mouse”, or “none”. When selecting “none”, skip the cell cycle score assignment. Default: “none”

#### Remove cell cycle

The user can remove the influence of the cell cycle phase on the dimensionality reduction results: “completely”, “partially”, or “do not remove”. When selecting “completely”, regress all signals associated with the cell cycle phase. For “partially”, regress only the differences in the cell cycle phase among proliferating cells; signals separating non-cycling and cycling cells will be maintained. When selecting “do not remove”, do not regress the signals associated with the cell cycle phase. Any selection will be ignored if the “Cell cycle gene set” input is not provided. Default: “do not remove”

#### Datasets metadata (optional)

If the selected sc analysis includes multiple aggregated datasets, each of them can be assigned to a separate group by one or multiple categories. This can be achieved by providing a TSV/CSV file with “library_id” as the first column and any number of additional columns with unique names, representing the desired grouping categories. To obtain a proper template for this file, download the “datasets_metadata.tsv” output from the “Files” tab of the selected “Single-cell Analysis with Filtered RNA-Seq Datasets” and add extra columns as needed.

#### Selected cell barcodes (optional)

Subsetting by cell barcodes can be used to identify sub-populations within the selected cluster(s) in the later steps. The user can obtain the list of barcodes by selecting the cells in the UCSC Cell Browser. Cells of interest are identified by their barcodes and stored in a TSV/CSV file with at least one column named “barcode”. All other columns, except for “barcode”, will be added to the sc metadata loaded from “Single-cell Analysis with Filtered RNA-Seq Datasets” and can be utilized in the current or future steps of analysis.

#### Number of highly variable genes

The number of highly variable genes is used in gene expression scaling, dataset integration, and dimensionality reduction. To keep only the biological signal and exclude possible noise, use the lower values for less heterogeneous datasets. The accepted range is from 500 to 5000 [41]. Default: 3000

#### Regress mitochondrial percentage

The user can regress the percentage of RNA reads mapped to mitochondrial genes as a confounding source of variation. Default: false

#### Regress genes

The user can regress expression of the selected genes as a confounding source of variation. Default: None

### 3 Single-Cell RNA-Seq Cluster Analysis

#### Target dimensionality

Target dimensionality is the number of principal components to be used in constructing the nearest-neighbor graph as part of the clustering algorithm. The accepted values range from 1 to 50. Default: 40

#### Clustering resolution

The resolution defines the “granularity” of the clustered data. Larger resolution values lead to more clusters. The optimal resolution often increases with the number of cells. For a dataset of 3,000 cells, a value within the 0.3-1.2 range usually returns good results. Default: 0.3

#### Find gene markers

The user can identify upregulated genes in each cluster compared to all other cells. The results include only genes that are expressed in at least 10% of the cells coming from either the current cluster or from all other clusters together. Genes with the log2FoldChange values smaller than 0.25 are excluded. The p-values are calculated with the Wilcoxon Rank Sum test and adjusted for multiple comparisons using the Bonferroni correction. Default: true

#### Genes of interest

A comma- or space-separated list of genes of interest can be used to visualize gene expression. Default: None

### 4 Single-Cell ATAC-Seq Dimensionality Reduction Analysis

#### Normalization method

The TF-IDF (Term Frequency–Inverse Document Frequency) normalization methods serve to correct for differences in cellular sequencing depth. Available options are: “log-tfidf” [9], “tf-logidf” [42], “logtf-logidf” [43] and “idf” [44]. Although each of these normalization methods produces reasonable results [43], we suggest using the default for Signac, which is the “log-tfidf” normalization method. Default: “log-tfidf”

#### Integration method

The integration methods serve to match shared cell types and states across experimental batches, donors, conditions or datasets: “signac” – uses cross-dataset pairs of cells that are in a matched biological state (“anchors”) to correct for technical differences; “harmony” – uses the Harmony algorithm to iteratively correct LSI (Latent Semantic Indexing) [45] embeddings; or “none” – does not run integration, merges datasets instead. Default: “signac”

#### Batch correction (harmony)

When “harmony” is selected as “Integration method”, batch effects are corrected on the basis of the provided factors. Specifically, “dataset” is used to integrate out the influence of the cells’ dataset of origin, while the factor “condition” is used to eliminate the influence of dataset grouping. Default: “dataset”

#### Target dimensionality

Target dimensionality is the number of dimensions to be used in LSI, datasets integration and UMAP projection. The accepted values range from 2 to 50. Default: 40

#### Datasets metadata (optional)

If the selected sc analysis includes multiple aggregated datasets, each of them can be assigned to a separate group by one or multiple categories. This can be achieved by providing a TSV/CSV file with “library_id” as the first column and any number of additional columns with unique names, representing the desired grouping categories. To obtain a proper template for this file, download the “datasets_metadata.tsv” output from the “Files” tab of the selected “Single-Cell Analysis with Filtered ATAC-Seq Datasets”, and add extra columns as needed.

#### Selected cell barcodes (optional)

Subsetting by cell barcodes can be used to identify sub-populations within the selected cluster(s) in the later steps. The user can obtain the list of barcodes by selecting the cells in the UCSC Cell Browser. Cells of interest are identified by their barcodes and stored in a TSV/CSV file with at least one column named “barcode”. All other columns, except for “barcode”, will be added to the sc metadata loaded from “Single-Cell Analysis with Filtered ATAC-Seq Datasets” and can be utilized in the current or future steps of analysis.

#### Minimum percentile of highly variable peaks

Set a minimum percentile for identifying the topmost common peaks as highly variable. For example, setting to 5 percent will use the top 95 percent most common among all cells peaks as highly variable. The selected peaks are then used for dataset integration, scaling and dimensionality reduction. Default: 0 (use all available peaks)

### 5 Single-Cell ATAC-Seq Cluster Analysis

#### Target dimensionality

Target dimensionality is the number of LSI components to be used in constructing the nearest-neighbor graph as part of the clustering algorithm. The accepted values range from 2 to 50. The first dimension is always excluded. Default: 40

#### Clustering resolution

The resolution defines the “granularity” of the clustered data. Larger values lead to more clusters. The optimal resolution often increases with the number of cells. Default: 0.3

#### Find peak markers

The user can identify differentially accessible peaks in each cluster compared to all other cells. The results include only peaks that are present in at least 5% of the cells coming from either the current cluster or from all other clusters together. Peaks with log2FoldChange values smaller than 0.25 are excluded. The p-values are calculated using the logistic regression framework and adjusted for multiple comparisons using the Bonferroni correction. Default: false

#### Genes of interest

A comma- or space-separated list of genes of interest can be used to generate ATAC fragment coverage plots. This is ignored if the “Cell Ranger ATAC or RNA+ATAC Sample (optional)” input is not provided. Default: None

### 6 Single-Cell WNN Cluster Analysis

#### Target RNA dimensionality

The target RNA dimensionality is the number of principal components to be used in constructing the weighted nearest-neighbor graph before clustering. The accepted values range from 1 to 50. Default: 40

#### Target ATAC dimensionality

The target ATAC dimensionality is the number of LSI dimensions to be used in constructing the weighted nearest-neighbor graph before clustering. The accepted values range from 2 to 50. The first dimension is always excluded. Default: 40

#### Clustering resolution

The resolution defines the “granularity” of the clustered data. Larger values lead to more clusters. The optimal resolution often increases with the number of cells. Default: 0.3

#### Find gene markers

The user can identify upregulated genes in each cluster compared to all other cells. The results include only genes that are expressed in at least 10% of the cells coming from either the current cluster or from all other clusters together. Genes with log2FoldChange values smaller than 0.25 are excluded. The p-values are calculated with the Wilcoxon Rank Sum test and adjusted for multiple comparisons using the Bonferroni correction. Default: true

#### Find peak markers

The user can identify differentially accessible peaks in each cluster compared to all other cells. The results include only peaks that are present in at least 5% of the cells coming from either the current cluster or from all other clusters together. Peaks with log2FoldChange values smaller than 0.25 are excluded. The p-values are calculated using the logistic regression framework and adjusted for multiple comparisons using the Bonferroni correction. Default: false

#### Genes of interest

A comma- or space-separated list of genes of interest can be used to visualize gene expression and to generate ATAC fragment coverage plots. This is ignored if the “Cell Ranger RNA+ATAC Sample (optional)” input is not provided. Default: None

### 7 Single-Cell Manual Cell Type Assignment

#### Cell types

Cell types are established using a TSV/CSV file with two columns named “cluster” and “celltype”. The first column includes the cluster numbers from the experiment selected in the “Single-cell Cluster Analysis” dropdown menu. The second column includes names to be assigned to each cluster. Depending the on the values of “Dimensionality reduction” and “Clustering resolution” parameters the same upstream experiment can be used for cell type assignment on the basis of scRNA-Seq, scATAC-Seq, or WNN clustering results.

#### Dimensionality reduction

Dimensionality reduction previously applied to the datasets. When combined with the “Clustering resolution” defines the cell groups for cluster names assignment. Available options are: “RNA”, “ATAC” and “WNN”. Default: “RNA”

#### Clustering resolution

Clustering resolution for the selected “Dimensionality reduction” is used to define clusters and assign cluster names.

#### Criteria to split every cluster by (optional)

The user can additionally split each cluster into several groups on the basis of the provided criteria. Available options are: “dataset”, “condition”, or “none”. Default: none

#### Find gene markers

The user can identify upregulated genes in each cell type compared to all other cells. The results include only genes that are expressed in at least 10% of the cells coming from either the current cell type or from all other cell types together. Genes with log2FoldChange values smaller than 0.25 are excluded. The p-values are calculated with the Wilcoxon Rank Sum test and adjusted for multiple comparisons using the Bonferroni correction. Default: true

#### Find peak markers

The user can identify differentially accessible peaks in each cell type compared to all other cells. The results include only peaks that are present in at least 5% of the cells coming from either the current cell type or from all other cell types together. Peaks with log2FoldChange values smaller than 0.25 are excluded. The p-values are calculated using the logistic regression framework and adjusted for multiple comparisons using the Bonferroni correction. Default: false

#### Genes of interest

A comma- or space-separated list of genes of interest can be used to visualize gene expression and to generate ATAC fragment coverage plots. This is ignored if the “Cell Ranger ATAC or RNA+ATAC Sample (optional)” input is not provided. Default: None

## Acknowledgments

This work was supported by the NHGRI grant R42HG011219 awarded to Datirium, LLC and the R01AI153442 awarded to Cincinnati Children’s Hospital Medical Center. We thank S. Hottinger for her editorial assistance. We are grateful to S. Potter, S. Korinfskaya, A. Shittu, J. Hellmann, and R. Player for their assistance with the interpretation of the analysis results which greatly enhanced the quality of this research. ChatGPT was used for language editing.

## Author contributions

A.B. and A.K. designed and developed SciDAP; M.K. and A.B. developed the CWL pipelines, analyzed data, and wrote the book chapter.

